# Automated plasmid design for marker-free genome editing in budding yeast

**DOI:** 10.1101/2024.11.12.623283

**Authors:** Lazar Stojković, Vojislav Gligorovski, Mahsa Geramimanesh, Marco Labagnara, Sahand Jamal Rahi

## Abstract

The ease of genome editing has contributed to the popularity of budding yeast as a model organism. However, the palette of selectable markers is in principle limited as most can only be used once. Some markers such as *URA3* and *TRP1* can be recycled through counterselection. This permits seamless genome modification with pop-in/pop-out (PIPO), in which a DNA construct first integrates in the genome and, subsequently, homologous regions recombine and excise undesired sequences. Popular approaches for creating such constructs use oligonucleotides and polymerase chain reaction (PCR). The drawbacks are that long oligonucleotides are unstable, can form secondary structures that interfere with PCR, cannot be regenerated in a typical biological laboratory, and are only widely available for lengths less than about 120, which limits the homology and efficiency that can be attained. With the rapid reduction in price, synthesizing custom DNA sequences in specific plasmid backbones has become an appealing alternative. For designing plasmids for seamless PIPO gene tagging or deletion, there are a number of factors to consider. To create only the shortest DNA sequences necessary, avoid errors in manual design, specify the amount of homology desired, and customize restriction sites, we created the computational tool PIPOline. Using it, we tested the ratios of homology that improve pop-out efficiency when targeting the genes *HTB2* or *WHI5*. We supply optimal PIPO plasmid sequences for tagging or deleting almost all S288C budding yeast open reading frames (ORFs). Finally, we demonstrate how the histone variant Htb2 marked with a red fluorescent protein can be used as a cell-cycle stage marker, alternative to superfolder GFP (sfGPF), reducing light toxicity. We expect PIPOline to streamline genome editing in budding yeast.

## Introduction

Budding yeast is an important model organism in genetics, molecular biology, and synthetic biology. Its genome is routinely modified in a highly targeted manner without specialized endonucleases; the homologous recombination machinery integrates the exogenous DNA into the yeast genome.^1^ Transformation can be performed with high efficiency but requires auxotrophic or antibiotic-resistance markers to screen for transformed cells^2–4^. To re-use markers, researchers utilize a two-step method, called pop-in/pop-out (PIPO), that removes the extraneous DNA after transformation, including the marker.^5–7^ Thus, in principle, an arbitrary number of modifications can be introduced in the genome. The method can be applied for deletions of open reading frames (ORFs) or seamless tagging with fluorescent proteins, for example (**Fig. 1**).

**Figure 1.**
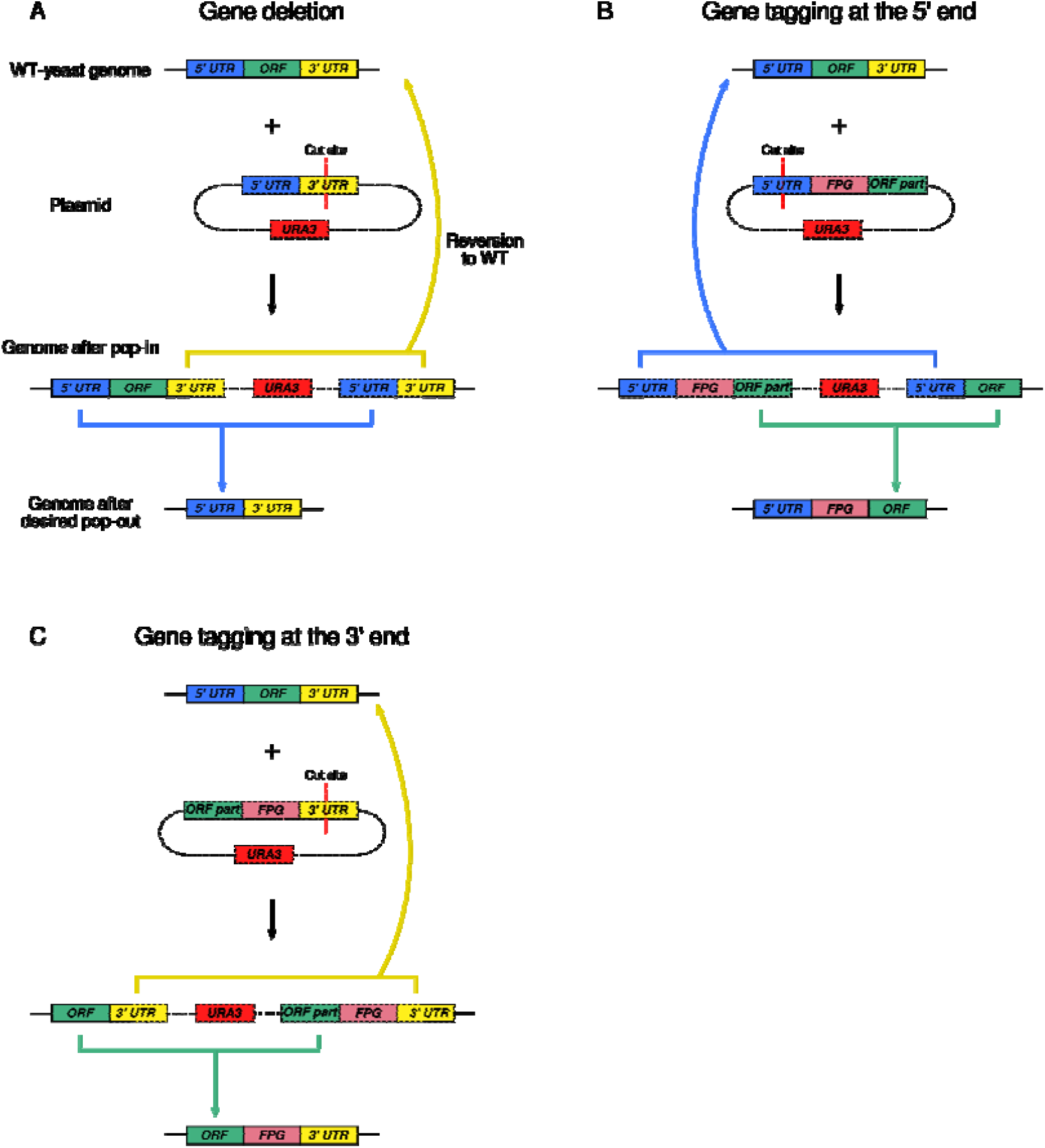
Gene editing using pop-in/pop-out (PIPO). A: Deletion of an open reading frame. Here, the pop-in homology comes from the 3’ UTR and pop-out homology from the 5’ UTR. In principle, the cut site for linearizing the plasmid could have been in a 5’ UTR sequence and the pop-out homology could come from the 3’ UTR (not shown). B: Genomic edit to tag the expressed protein at the N-terminus. In this illustration, the pop-in homology comes from the 5’ UTR and the pop-out homology from the ORF. In principle, the cut site for linearizing the plasmid could have been in a sequence from the ORF and the pop-out homology could come from the 5’ UTR (not shown). C: Genomic edit to tag the expressed protein at the C-terminus. Here, the pop-in homology comes from the 3’ UTR and the pop-out homology from the ORF. The roles of the 3’ UTR and ORF for pop-in and pop-out homology could have been reversed (not shown). A, B, C: PIPO is performed using a custom-designed plasmid that allows seamless two-step genome editing. In the first step, the PIPO plasmid is integrated and cells are selected for the *URA3* marker. In the second step, cells are selected for the loss of *URA3*. Depending on the DNA sequences that recombine, pop-out can either lead to reversion to the wild-type yeast genome (curved arrows pointing up) or to the desired genome edit (colored arrows pointing down). A, B, C: *FPG* – fluorescent protein gene, *UTR* – untranslated region, *ORF* – open reading frame.

PIPO involves initial positive selection followed by negative selection (counterselection). Both of these steps select for cells in which recombination events occur: In the pop-in step, the sequence of interest integrates into the yeast genome, thereby introducing the selectable marker. In the pop-out step, cells that have lost the marker are selected. Critically, depending on the recombination site, these cells may revert to the wild-type (WT) genome (curved arrows pointing up in **Fig. 1**), which is undesired, or result in the desired genome modification (colored arrows pointing down in **Fig. 1**).

Marker genes that can be both selected for and against in budding yeast are in the lysine^8^, tryptophan^9^, or uracil^10^ biosynthesis pathways. Among them, the *URA3* gene is often used due to its short length (267 amino acids) and the low spontaneous reversion frequency of the genomic *ura3Δ* copy. *URA3* is selectable on uracil-deficient media and counterselectable on media containing 5-fluoroorotic acid (5-FOA)^10^. The experimental process is depicted in **Fig. 2**.

**Figure 2.**
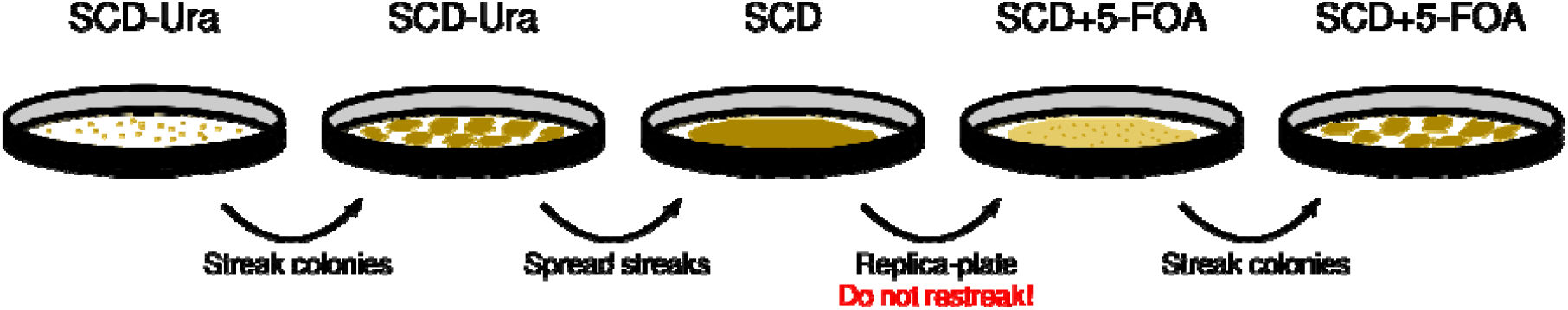
Experimental steps in PIPO. Cells transformed with the PIPO construct are selected on synthetic complete media plates lacking uracil (SCD-Ura). Transformant colonies are then restreaked on fresh SCD-Ura plates. One or multiple of these patches are then spread onto non-selective SCD plates (one SCD plate for each original transformant colony). After one day, cells are transferred to an SCD plate containing 5-FOA (SCD+FOA) by replica plating. After about 4 days, single colonies from the SCD+5-FOA plate are restreaked onto a fresh SCD+FOA plate.

An advance in the design of cassettes for seamless genome modifications was to directly use PCR-generated DNA sequences to replace a genomic sequence of interest^11,12^, used for example in the creation of the yeast knockout collection^13^. In doing so, researchers skip the plasmid construction step. This has led to the design of several innovative strategies, such as “delitto perfetto”^14^ and 50:50^15^. However, PCR-based transformation is limited in the amount of homology that can be created for genomic integration. Commercially available oligonucleotides are typically shorter than 120 nucleotides. The efficiency of such transformations is dependent on the strain background.^12^ In addition, oligonucleotides are less stable than plasmids and cannot be easily regenerated in typical biological laboratories. This led us to focus on plasmid-based PIPO, in part, since the prices for DNA synthesis and cloning have dropped sharply.

There are a number of parameters that have to be simultaneously optimized for plasmid-based PIPO, making manual design tedious:

- sufficient, user-specified minimum homology for plasmid integration (pop-in);
- specific amount of homology for plasmid pop-out;
- minimal length of the DNA sequence to be synthesized;
- uniqueness of restriction site for plasmid integration; and
- restriction sites for subcloning different tags such as fluorescent protein genes.

These criteria can be formalized for a computational pipeline. Here, we present PIPOline (<u>P</u>op-in/<u>p</u>op-<u>o</u>ut pipe<u>line</u>), a Python-based algorithm for the design of plasmids for PIPO-based genome editing. Additionally, we use PIPOline to show how the probability of the desired pop-out can be fine-tuned through sequence length optimization. Finally, labeling the histone variant Htb2, we demonstrate how cell-cycle phases can be observed using the fast-folding red fluorescent protein ymScarletI, alternative to sfGFP.

## Results

### PIPOline: pipeline for the design of pop-in/pop-out plasmids

We designed PIPOline for two types of genomic modification: ORF deletion or protein tagging at the N- or C-terminus. The inputs to the PIPOline program are listed in **Table 1**; the names of the corresponding command line parameters are supplied in **Supplementary Table 1**. PIPOline can in principle design a PIPO plasmid for any vector backbone. To easily eliminate *URA3* revertants during pop-in and pop-out steps, we created a dually marked backbone, pETURALEU, available from the Addgene plasmid repository. In addition to the *URA3* auxotrophic marker, the plasmid contains the *LEU2* gene. In a strain with a *LEU2* loss-of-function mutation, *URA3* revertants can be easily distinguished from cells having undergone successful pop-in or pop-out by additionally testing for leucine auxotrophy. (In contrast to pETURALEU, the pLS406 shuttle vector^16^ only has a *URA3* marker and was miniaturized and depleted of most restriction sites outside the MCS to improve the efficiency of complex insert engineering such as for PIPO.)

**Table 1.**
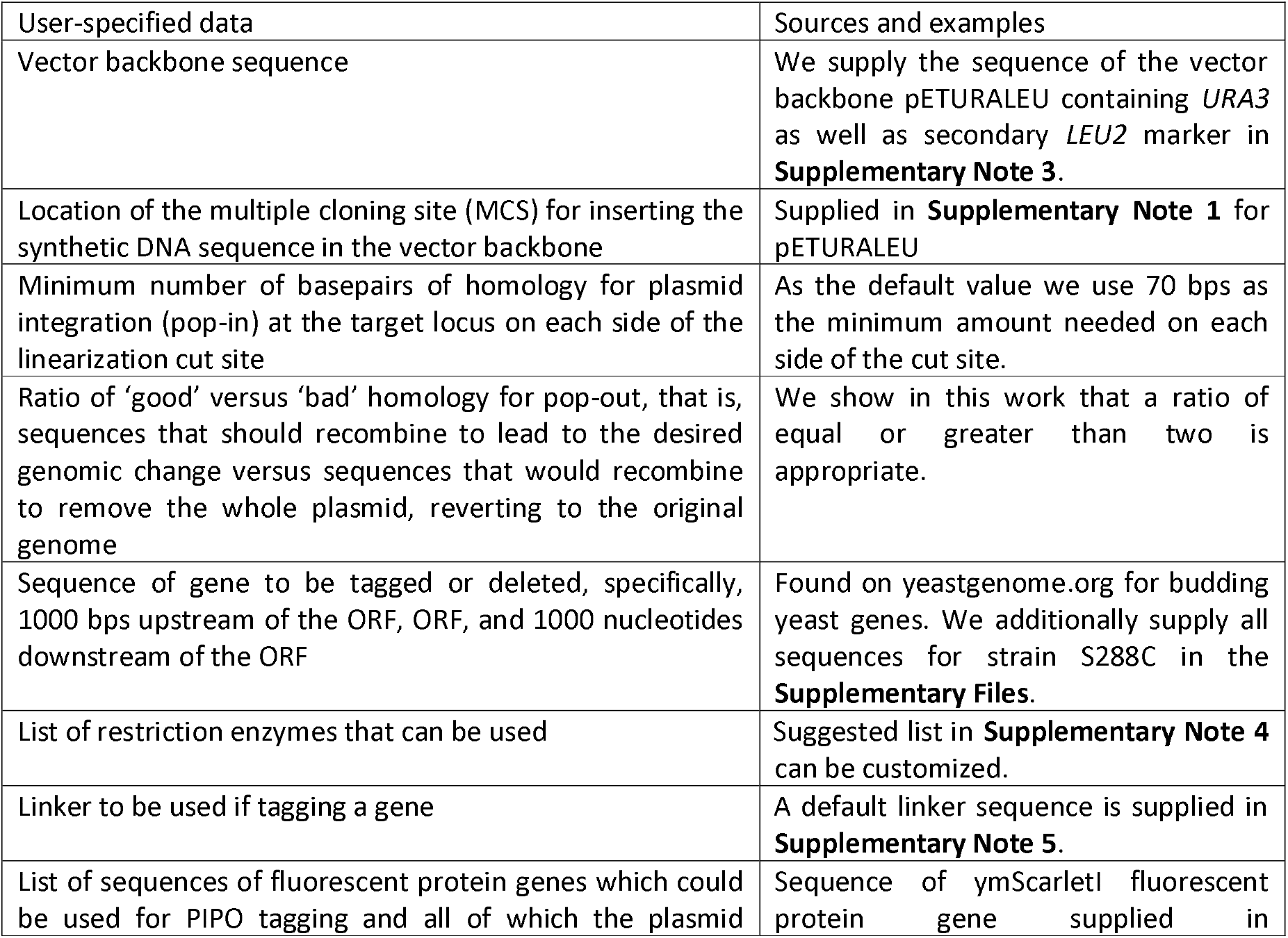
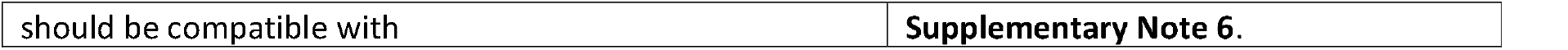
Inputs to PIPOline program.

| User-specified data | Sources and examples |
| --- | --- |
| Vector backbone sequence | We supply the sequence of the vector backbone pETURALEU containing *URA3* as well as secondary *LEU2* marker in **Supplementary Note 3**. |
| Location of the multiple cloning site (MCS) for inserting the synthetic DNA sequence in the vector backbone | Supplied in **Supplementary Note 1** for pETURALEU |
| Minimum number of basepairs of homology for plasmid integration (pop-in) at the target locus on each side of the linearization cut site | As the default value we use 70 bps as the minimum amount needed on each side of the cut site. |
| Ratio of ‘good’ versus ‘bad’ homology for pop-out, that is, sequences that should recombine to lead to the desired genomic change versus sequences that would recombine to remove the whole plasmid, reverting to the original genome | We show in this work that a ratio of equal or greater than two is appropriate. |
| Sequence of gene to be tagged or deleted, specifically, 1000 bps upstream of the ORF, ORF, and 1000 nucleotides downstream of the ORF | Found on yeastgenome.org for budding yeast genes. We additionally supply all sequences for strain S288C in the **Supplementary Files**. |
| List of restriction enzymes that can be used | Suggested list in **Supplementary Note 4** can be customized. |
| Linker to be used if tagging a gene | A default linker sequence is supplied in **Supplementary Note 5**. |
| List of sequences of fluorescent protein genes which could be used for PIPO tagging and all of which the plasmid | Sequence of ymScarletI fluorescent protein gene supplied in |

The steps in PIPOline are enumerated below and depicted in **Fig. 3**. In summary, dependent on the genomic editing task with PIPOline, two sequences from different regions of a gene of interest (GOI) need to be included in the PIPO plasmid, taken from the 5’ UTR, ORF, or 3’ UTR. The algorithm loops through candidate restriction sites in the two regions that may be used for linearizing the final plasmid. The pipeline evaluates whether the PIPO plasmid can be built based on the candidate restriction site according to a specific set of rules. If not, the program notifies the user about the reason why a candidate linearization cut site has to be disregarded. Otherwise, the algorithm outputs the sequence to be synthesized for each linearization cut site, which can be synthesized and cloned into the specified vector backbone:

**Figure 3:**
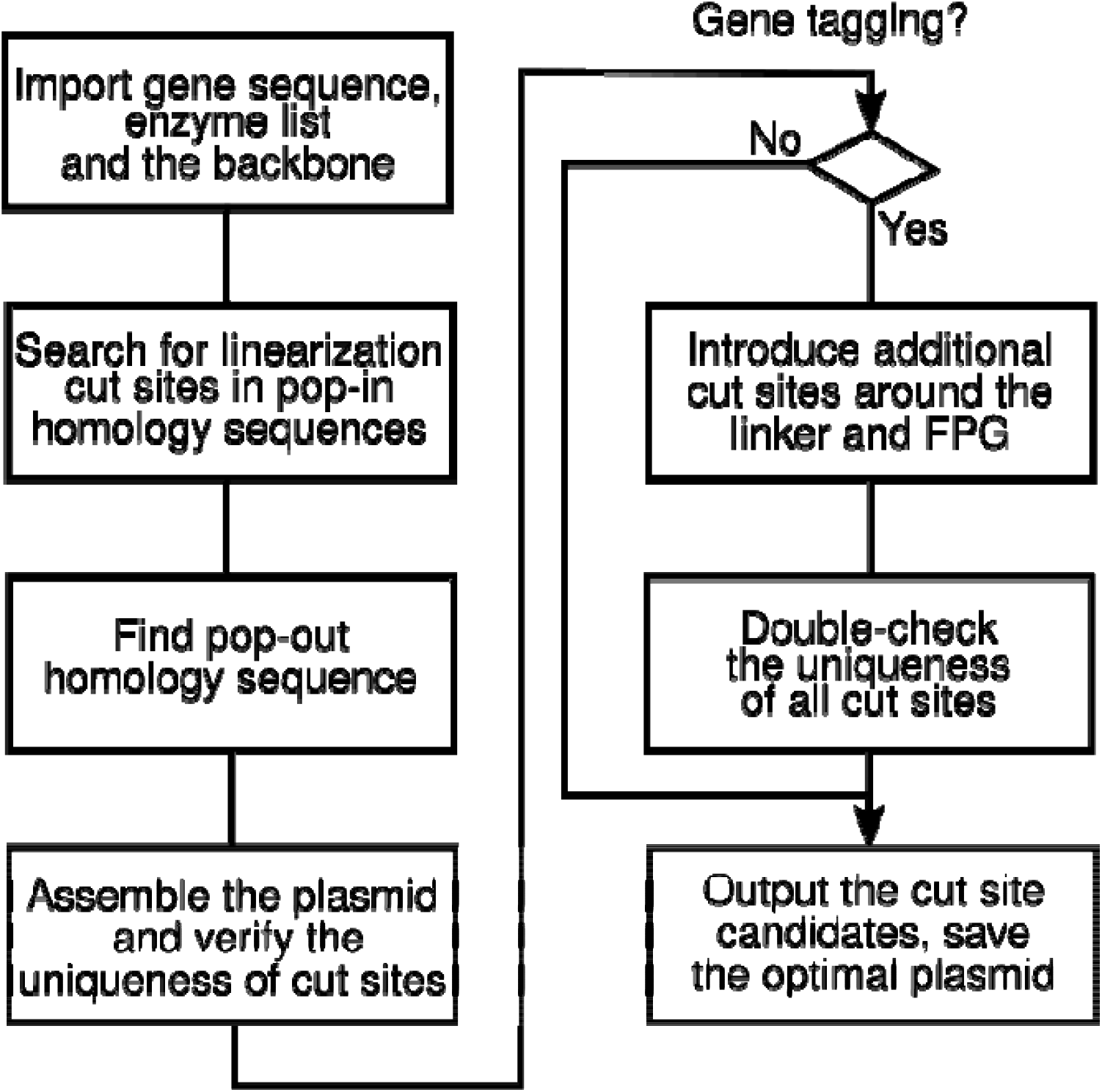
PIPOline process diagram.

1. For tagging or deleting ORFs, two regions of the gene of interest (GOI) are used to create homology for pop-in and pop-out, respectively: 1) 5’ UTR and ORF for tagging the ORF at the 5’ terminus, 2) 3’ UTR and ORF for tagging the ORF at the 3’ end, or 3) 5’ UTR and 3’ UTR for deleting the ORF. To determine the 5’ UTR, ORF, and 3’ UTR, the PIPOline algorithm assumes that the user-supplied GOI sequence consists of 1000 bps downstream of the ORF, followed by the ORF, and 1000 bps upstream of the ORF. Parts of both regions are used in the PIPO plasmid, one for pop-in and the other for pop-out, as determined by the following steps. The code performs two checks: i) for Start and Stop codons at the expected positions and ii) whether there is overlap with another ORF (based on similarity to overlapping ORFs in budding yeast strain S228C), and alerts the user. For each budding yeast gene, the required DNA sequence can be readily downloaded from the yeastgenome.org database^17^. We additionally supply all such sequences for budding yeast strain S288C in the **Supplementary Files**.
2. One of the two regions will be used for integrating the plasmid into the genome (“pop-in homology”, boxes containing cut sites in **Fig. 1**). A sequence from this region will be included in the PIPO plasmid and contains the cut site used to linearize the plasmid with restriction enzymes prior to transformation. To identify the linearization cut site and the pop-in homology, PIPOline searches in both regions for unique restriction sites corresponding to the user’s preferred restriction enzymes (Table 1). For each candidate linearization cut site, PIPOline measures the size of the region on both sides of the cut site. These sequences must be longer than a user-specified minimum. If cutting the plasmid at the candidate cut site results in less than the specified minimum homology, the linearization cut site is not considered further by the program. For example, the cut site in **Fig. 1** A has to be sufficiently far from the 5’ UTR sequence to its left. As pop-in homology, PIPOline takes the whole sequence between the cut site and the neighboring region as well as a sequence of length corresponding to the minimum homology on the other side of the cut site.
3. The region which is not used for linearizing the plasmid and integration is required for the counterselection step (“pop-out homology”, e.g., blue box in **Fig. 1 A**). The user-supplied parameter R_homo_ specifies the ratio of pop-out to pop-in homology. PIPOline checks whether a sufficiently long sequence is available in the second region (i.e., region not used for pop-in homology) and uses it for pop-out homology. If a sufficiently long sequence is not available, the maximal available sequence is taken for pop-out homology, and a warning message is associated with this linearization cut site and construct.
4. The sequences of pop-in and pop-out homology as well as a linker and fluorescent-protein gene (FPG) for tagging the GOI are assembled for a candidate pop-in/pop-out construct.
5. Unique restriction sites for cloning the insert into the multiple cloning site of the vector backbone, maintaining the uniqueness of the linearization restriction site, are identified. If this is not possible, the linearization cut site is disregarded and a message describing the error is displayed.
6. For ORF tagging, PIPOline searches for additional cut sites to be added around the linker and around the fluorescent protein gene sequence. This allows for straightforward exchange of these pieces in the future. The program uses a user-defined list of restriction enzymes with 6 bp long cut sites and verifies that they do not introduce a stop codon.
7. PIPOline outputs the list of all linearization cut site candidates. Cut sites which cannot be used for constructing a PIPO plasmid respecting the above rules are listed as well, together with the reason for discarding them. If a linearization cut site can be used for a PIPO plasmid, the program outputs the sequences that need to be synthesized, the restriction sites for sub-cloning, and the total length of the insert DNA. The linearization cut site that meets all criteria and requires the minimal amount of DNA to be synthesized is considered to be optimal, and the corresponding assembled plasmid sequence is saved as a FASTA file.

We tested the algorithm with all 6039 budding yeast strain S288C genes identified at yeastgenome.org for ORF tagging or deletion with R_homo_ = 2 (see below). PIPOline could find sequences satisfying the above criteria for 99.42% of the plasmids (18012 out of 18117) needed to tag or delete each gene. Further statistics regarding the plasmids are supplied in **Supplementary Figs. 1** and **2**. In the **Supplementary Files**, we supply all plasmid sequences for deletion or tagging; the desired tag sequence, with potential duplicate restriction sites removed by switching synonymous codons, has to be inserted by the user.

PIPOline is available as a command-line tool (**Code availability**) and has been tested on Linux Ubuntu 21.04 and Windows 10 operating systems. Examples of calls for different genome editing tasks are given in **Supplementary Note 2**. The tool can also be run in an online graphical user interface available at https://colab.research.google.com/drive/12hle7b94_j6DbAeaJFaCH6jbU1yM1f_Z.

### Testing PIPOline and maximizing successful pop-outs by tuning R_homo_

In the pop-out step, the pop-in and pop-out homology of the integrated PIPO plasmid can recombine with homologous sequences in the genome, resulting either in the desired genomic change or reversion to the wild-type sequence (colored arrows in **Fig. 1**). Therefore, the colonies that emerge after counterselection have to be screened for the desired pop-out. We investigated whether tuning the ratio of lengths of pop-in and pop-out homology could favor one recombination event over the other. This would guide plasmid design for efficient pop-out and reduce the manual screening burden. Such a strategy has been considered before^1^ and would plausibly work due to the dependence of recombination frequencies on the lengths of homologous sequences^18–21^.

To test the idea that competition between the two spontaneous recombination outcomes is decided by the ratio of lengths of homologous sequences, we used PIPOline to construct a series of plasmids that would C-terminally tag Htb2 or Whi5. In each series, the pop-in homology sequence was kept constant, while the pop-out homology length was varied. Thus, R_homo_ ranged between 0.5 and 4 for Whi5 tagging and between 0.5 and 2 for Htb2 tagging. (Extending R_homo_ to 4 for Htb2 tagging was not possible since it would have led to a non-unique linearization cut site.)

We created the PIPOline inserts with the different R_homo_ values in the pETURALEU plasmid backbone. (Alternatively, these inserts could be obtained from commercial gene synthesis vendors, cloned in the desired vector backbone.) We linearized plasmids with the enzymes chosen by PIPOline and performed a standard LiAc chemical transformation with 0.5 ug of DNA. Transformants were selected on synthetic complete glucose agar plates without uracil (SCD-Ura) (**Fig. 2**). Starting with 5 mL of exponentially growing culture, we typically obtain up to 30 transformants. Colonies were restreaked as patches onto fresh SCD-Ura plates to minimize the chance that untransformed cells could be transferred to the next steps. To allow spontaneous recombination events to pop out the undesired sequences, we then streaked out 2-3 of the patches onto non-selective growth medium plates. For the next step, we found it important to replica plate from the non-selective growth medium plate onto a 5-FOA plate and not streak a sample from the non-selective medium plate (**Methods**). We hypothesize the reason to be that many genomic edits result in a fitness penalty,^22,23^ which would increase the relative number of cells with revertant pop-outs versus desirable pop-outs if cells are mixed (e.g. restreaked). Replica plating from non-selective to counter-selective plates should not change the relative number of colonies.

When pop-out occurs in the desired direction, the *URA3* and *LEU2* markers are removed from the genome, leaving the fluorescent protein at the ORF 3’ terminus. On the other hand, if the whole plasmid pops out from the genome, the gene remains unlabeled. We scored the type of resulting pop out using two different strategies: fluorescence microscopy for *HTB2* tagging and PCR for *WHI5* tagging.

As expected, the probability of the desired pop-out for equal lengths of pop-in and pop-out homology sequences (R_homo_ = 1) was close to half (55% for *HTB2* and 39% for *WHI5*) (**Fig. 4**). Interestingly, for both *HTB2* and WHI5 we observed that the efficiency of the desired pop-out was tunable across a range of probabilities by modulating R_homo._ Reducing R_homo_ to 0.5 made the desired pop-outs very improbable (8% for *HTB2* and 2% for *WHI5*). On the other hand, increasing it to 2 led to 79% or 89% of the colonies having the desired pop-out, when tagging *HTB2* or WHI5, respectively (**Fig. 4**). Increasing R_homo_ above 2 in the case of *WHI5* had no apparent effect on pop-out efficiency.

**Figure 4.**
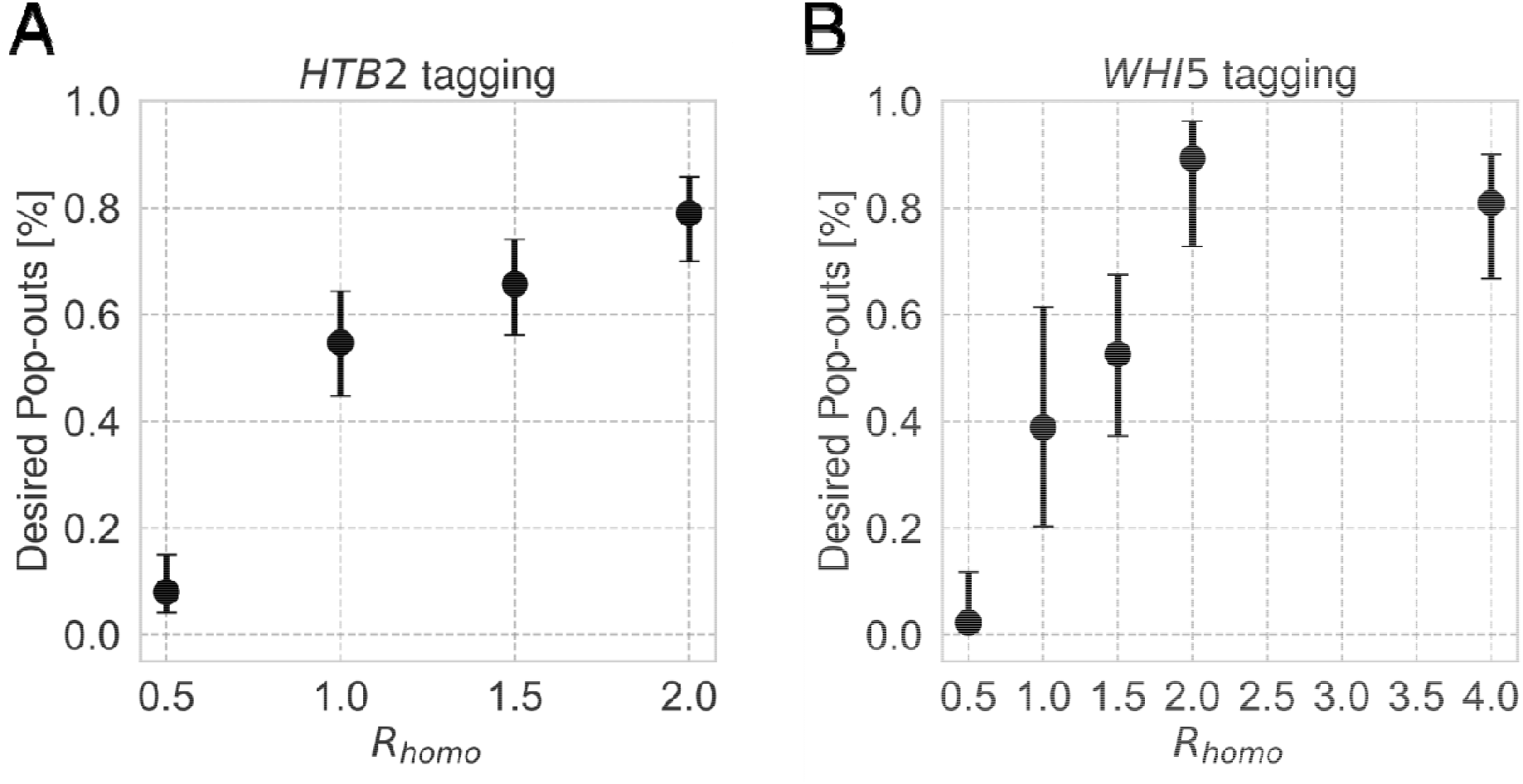
Increasing the ratio of lengths of pop-out versus pop-in homology (R_homo_) leads to a higher probability of the desired pop-out when tagging either A: *HTB2* or B: *WHI5* in budding yeast. Error bars show 95% confidence intervals calculated using the Wilson score method. The numbers of analyzed individual colonies are shown in Supplementary Table 3.

The pop-out efficiency was scored using at least two different transformed (pop-in) colonies. We observed that in some cases, some colonies transformed with the PIPO plasmid never popped out the plasmid in the desired manner despite a relatively large R_homo_ (1.5 and 4). We did not identify the reason for this. We exclude such colonies from the statistics presented in **Fig. 4** based on the very different pop-out efficiency compared to other colonies transformed with the same plasmid. This observation implies that it is best to use R_homo_ of about 2 or larger and also use more than one transformant when counterselecting for pop-out (**Discussion**).

Based on these results, we applied PIPOline to tagging 7 genes and deleting 8 genes. We ordered the insert corresponding to the PIPOline output to be synthesized and cloned into the pETURALEU vector backbone. While we did not quantify the number of pop-out colonies screened, we readily succeeded in tagging *CDC5, CLB2, CLB5, DNL4, NEJ1, SIC1*, and *SPC42* and in deleting *CLB3, CLB4, CLB5, CLB6, HEM14, HMX1, MRE11*, and *YMR315W*. The specific inserts and plasmid sequences we used are supplied in the **Supplementary Files**.

### Resolving cell-cycle phases using ymScarletI-marked histones

Finally, we wondered whether an Htb2-ymScarletI fusion created using PIPOline allows monitoring of histone accumulation in S phase. Researchers have previously shown that the pulse-like pattern of histone transcription during S-phase^24,25^ can be tracked using sfGFP^26,27^ but not with the relatively slow-folding mCherry^26^. However, monitoring cellular processes using red fluorescent proteins produces less phototoxicity than with GFP. This is particularly important when frequent imaging is needed as in the case of dynamic cell cycle-related events.

The maturation time of ymScarletI in budding yeast is 12 min^28^, which is relatively close to that of sfGFP (7 min^27^) compared to mCherry (52 min^27^). To test whether ymScarletI suffices to observe histone accumulation in single cells, we performed timelapse fluorescent microscopy. To correlate fluorescence levels more easily with the cell-cycle stage, we scaled the time axis: We assigned the moment in which the nuclei split into mother and daughter cells (anaphase) to timepoint zero, and followed the daughter cell through budding until rebudding, assigned to timepoint one.

Histone synthesis occurs in S phase after cells have passed Start and committed to budding.^29^ Hence, we expected to see accumulating fluorescence around the budding time (dashed lines in **Fig. 5 A**). Indeed, shortly after budding, fluorescence levels started to rise and in the subsequent anaphase dropped to about half of the maximal value. Note that in the black curve which represents the average of the normalized single-cell timecourses in **Fig. 5**, the peak is smeared out due to variability in the lengths of individual cell cycles. The region with flat fluorescence corresponded to G1 phase, which is substantially longer in daughter cells compared to mother cells (compare single-cell traces around timepoints 0 and 0.7). These results illustrate how two critical points of the cell cycle, G1-to-S transition and anaphase, can be visualized by a single red-fluorescent marker.

**Figure 5.**
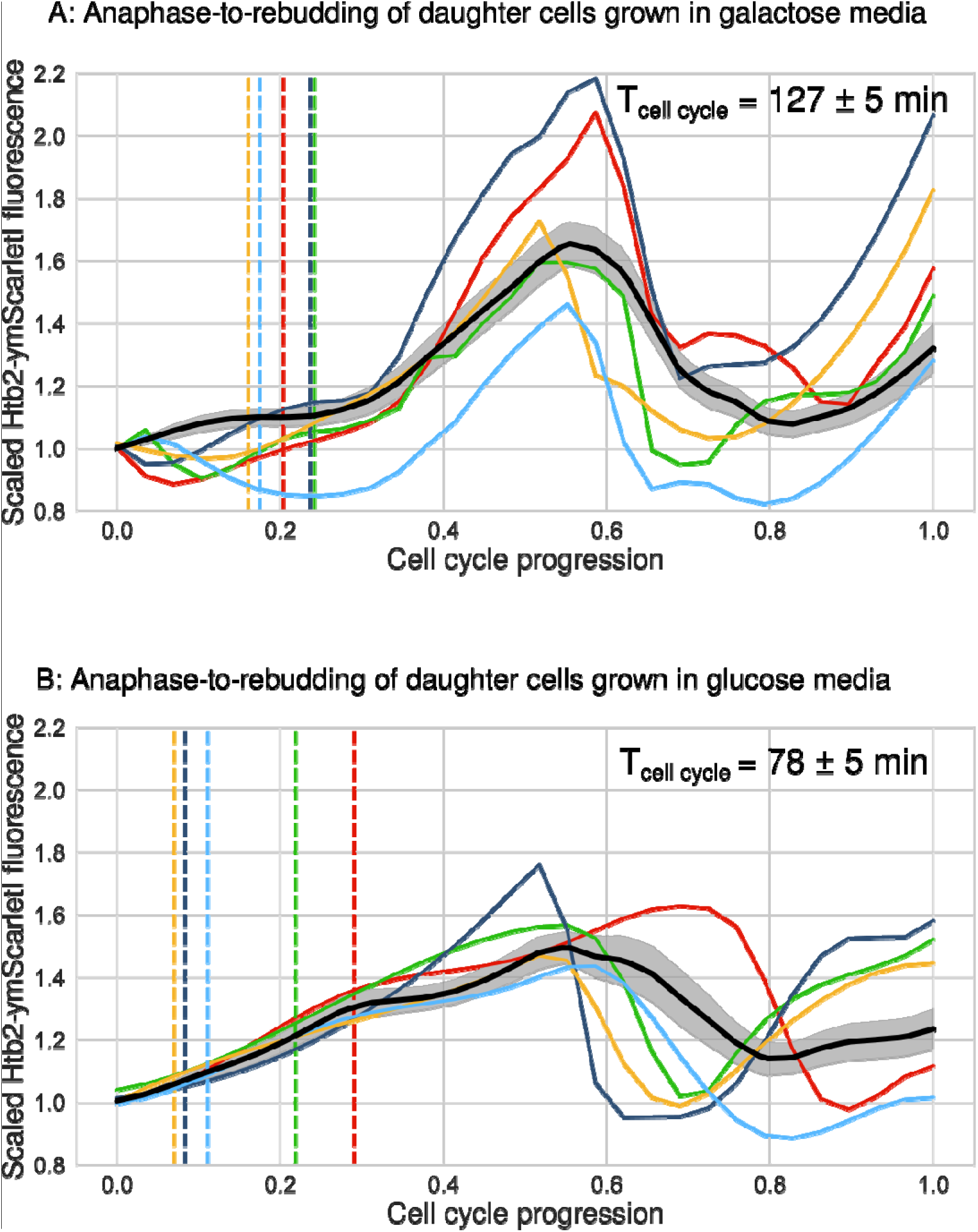
Htb2-ymScarletI dynamics in single daughter cells, from mother-daughter anaphase at time 0 to daughter rebudding at time 1. Cells were grown in synthetic complete media supplemented with A: 3% w/v galactose or B: 2% w/v glucose. Colored traces illustrate single-cell timecourses while population averages are denoted in black. Shown are total fluorescent intensities, scaled by their first timepoint. To accommodate different cell-cycle lengths, as in a previous study involving histone synthesis tracking ^27^, we normalized the time axis. Dashed lines show budding events, which in slowly cycling cells (A) precede histone accumulation. In media supplemented with galactose (A), the average budding-to-rebudding time was 127 ± 5 min (mean ± standard error of the mean), while in media with glucose (B) it was 78 ± 5 min. Activation of histone synthesis in S phase can be observed in the slow cel cycles (A) but not in the fast ones (B). The numbers of analyzed cells were A: 29 and B: 13. The standard errors of the mean (shaded area around the thick black line in both of the panels) showed that this number of cycles was sufficient to estimate the dynamics precisely.

In this and previously published experiments involving Htb2-sfGFP, cells were grown in galactose media, glucose-limited media^26^, or minimal media^27^, resulting in a long cell-cycle period (127 min in our experiment and 145 min in ref. ^26^). However, cells are often instead grown in rich media supplemented with 2% glucose, where the average cell cycle period is about 80 min^30^. Under such conditions, the transition from G1 to S phase might not be as obvious. Indeed, for cells grown in glucose-rich media (**Fig. 5 B**), the G1 period was less clear with fluorescence increasing more gradually. The time between anaphase to daughter budding was about 32 ± 3 min (mean ± standard error of the mean). Hence, ymScarlet may mature too slowly to observe histone dynamics in budding yeast cells growing in glucose-rich medium.

In summary, we conclude that HTB2-ymScarletI maturation allows visualization of progression through Start and anaphase in relatively slow cell cycles in a manner comparable to HTB2-sfGFP, the only other such marker available so far.

## Discussion

We present a pipeline for the automated design of plasmids for PIPO for tagging or deleting genes in budding yeast. PIPOline is customizable through user-defined parameters. It outputs a variety of possible strategies for a desired genomic edit and identifies the smallest insert that needs to be synthesized. PIPOline assumes the first and last 1000 bps in the GOI file are 5’ UTR and 3’ UTR. Hence, designing plasmids for intergenic labeling using PIPOline is possible by adjusting the input file.

We experimentally mapped the probability of the desired recombination event depending on the ratio of pop-out and pop-in homology sequence lengths. This showed that having the pop-out homology twice as long compared to the pop-in homology leads to a large portion (>79%) of the popped-out colonies having the desired genomic edit. However, some colonies transformed with the PIPO plasmid did not yield any desirable pop-outs. Hence, the strategy that we propose is to use a R_homo_ of at least two and pop out from at least two different transformed colonies. In our tests, this sufficed for finding at least one desired genotype.

PIPOline is well-suited for genomic modifications in an era where commercial vendors offer gene synthesis and cloning into custom vectors at prices that are relatively affordable for typical laboratories. Using, for example, the doubly *URA3*- and *LEU2*-marked pETURALEU plasmid, only the minimal PIPOline-designed insert needs to be synthesized and cloned into the vector backbone. There are a number of advantages over oligonucleotide-based PIPO approaches: Plasmids are easier to reproduce, reuse, share, and store long term. Because commercial oligonucleotides are limited in length to about 120 nts, the amount of homology that they can encode is limited to about 50 bps for insertion and 50 bps for pop-out, near the minimum limit for the homologous recombination machinery^18–21^. Furthermore, the PCR step to create the PIPO construct becomes unreliable with long oligonucleotides. Finally, in the market relevant to our laboratory, the price per base pair for gene synthesis is lower than per nucleotide for oligonucleotides.

There are several similarities and differences between PIPO and CRISPR-Cas9-based scarless editing. Here, we assume that all of the cloning steps for CRISPR-Cas9 editing are performed by commercial gene synthesis service providers as well, eliminating differences in the amount of labor involved in preparing DNA constructs. Cas9 needs a single guide RNA (sgRNA) to target a specific genomic locus. The design of the sgRNA must respect the need for a protospacer-adjacent motif (PAM) and minimize off-target cleavage^31^, making it common to use computational tools for this step. After inducing a cut at the target site, a co-transformed template DNA is used by the cell’s DNA repair machinery to introduce the desired genomic change^31^. The DNA constructs should be designed in such a way that in this step, the sequences recognized by Cas9 are eliminated to prevent re-cutting. Finally, the plasmids that have been introduced are counter-selected or lost during proliferation in non-selective growth media. The number of steps and time needed for the two methods is roughly comparable, one transformation and one counterselection. Instead, we see the main difference in the troubleshooting that may be needed: The first step in PIPO, transformation with an integrating plasmid, is generally efficient and is performed routinely in yeast laboratories. Failure modes and counter-measures are well-known. The factors important for the second step in PIPO, pop-out by spontaneous recombination, have been characterized in our work presented here. On the other hand, what steps to take when CRISPR-Cas9 genome editing fails is much less well known in typical yeast laboratories. This difference in the prevalence of expertise in the two techniques will diminish if CRISPR technology is more broadly adopted in yeast laboratories. Regardless, PIPOline fills a gap in making the currently most prevalent strategy for genome editing in budding yeast, which does not rely on endonucleases, more efficient.

## Methods

### Yeast strain construction

Transformations of budding yeast were performed using the high-efficiency LiAc/DNA carrier/PEG (polyethylene glycol) protocol^32^. All strains are derivatives of W303 (*leu2-3,112 trp1-1 can1-100 ura3-1 ade2-1 his3-11,15*). The list of strains used is supplied in **Supplementary Table 1**.

### Media

We used synthetic complete media^33^ (SC) with or without uracil (Ura) and supplemented with 2% (w/v) glucose (D) or 3% (w/v) galactose (G). For preparing solid SC plates, we used 2% agar. For experiments with 5-FOA (SCD+5-FOA), we used the standard recipe with 0.1% 5-FOA^10,34^.

### Plasmid construction

Plasmids designed by PIPOline were assembled using restriction enzyme cloning and T4 ligase (NEB). The regions homologous to the yeast genome were amplified using high-fidelity Phusion polymerase (Thermo Fisher Scientific) with genomic DNA as a template. Yeast codon-optimized *mScarletI* (*ymScarletI*) was adapted from ref. ^35^ by removing cut sites of frequently used restriction enzymes and was synthesized by GenScript. Plasmids were propagated in XL10 *E. coli* in LB medium supplemented with ampicillin. The list of plasmids used is supplied in the **Supplementary Table 2**.

### Pop-in/pop-out

Cells were transformed with the pETURALEU plasmid containing the insert used to modify the locus of interest. After 3 days, isolated colonies were restreaked onto fresh SCD-Ura plates. Upon confirming that the plasmid integrated as expected into the yeast genome by PCR and selection on –Leu plates, we proceeded to the pop-out step. First, cells were grown on non-selective media (SCD) plates to enable the survival of the colonies in which the *URA3* marker was lost. To spread cells onto SCD plates, single colony streaks from SCD-Ura were briefly resuspended in 100 uL of liquid SCD medium and then evenly spread onto agar plates. After 24 h, cells were replicated on SCD+5-FOA plates to select for cells that did not contain the *URA3* marker. Cells were grown for 4 days on FOA after which single colonies were picked and regrown on SCD+5-FOA plates before the screen. For a graphical summary, see **Fig. 2**.

### Fluorescence microscopy

Recordings were made using Nikon Ti-2E microscope equipped with a 20x objective and a Hamamatsu Orca-Flash4.0 camera. For scoring the pop-out efficiency, the strains were grown in 96 well-plates in synthetic complete medium. The strains were scored based on the intensity of the nuclear foci in the red fluorescence channel with 50 ms exposure time. For running timecourse experiments, cells were grown in CellAsic microfluidic chips and recorded using the same microscope with 63x objective. For the timelapse recordings, images were taken every 5 minutes.

## Data analysis

For processing single-cell microscopy experiments, cells were segmented and tracked using YeaZ^36^, available at https://github.com/rahi-lab/YeaZ-GUI. For analyzing single-cell timecourses we used a custom-based code written in Python 3.11.3.

Individual cell cycles shown in **Fig. 5** had variable duration. To make their dynamics comparable, we normalized the time axis so that timepoint 0 corresponds to mother-daughter anaphase and 1 to the budding of the daughter cell. To calculate the average timecourse, single-cell curves were interpolated at 30 timepoints and smoothed using spline interpolation (splrep function from scipy 1.10.1 Python package with smoothing factor 0.02).

## Data availability

pETURALEU is deposited with Addgene (ID: 223207). The data are available from the corresponding author upon request.

## Code availability

The PIPOline code, tested in Python 3.11.3, is available at https://github.com/rahi-lab/PIPOline. A web application is also available at: https://colab.research.google.com/drive/12hle7b94_j6DbAeaJFaCH6jbU1yM1f_Z.

## Contributions

LS, VG, and SJR designed PIPOline and wrote the code. ML tested the code and provided feedback. VG constructed the plasmids and strains. MG and VG evaluated optimal homology ratios. VG and SJR wrote the manuscript. SJR supervised the project and acquired funding.

## Acknowledgements

We thank Dr Enrico Tenaglia for the help with molecular cloning. We acknowledge support for VG, ML, and SJR provided by SNSF grants CRSK-3_190526, 310030_204938, and CRSK-3_221036 awarded to SJR.

## Supplementary information

### Supplementary Figures

**Supplementary Figure 1.**
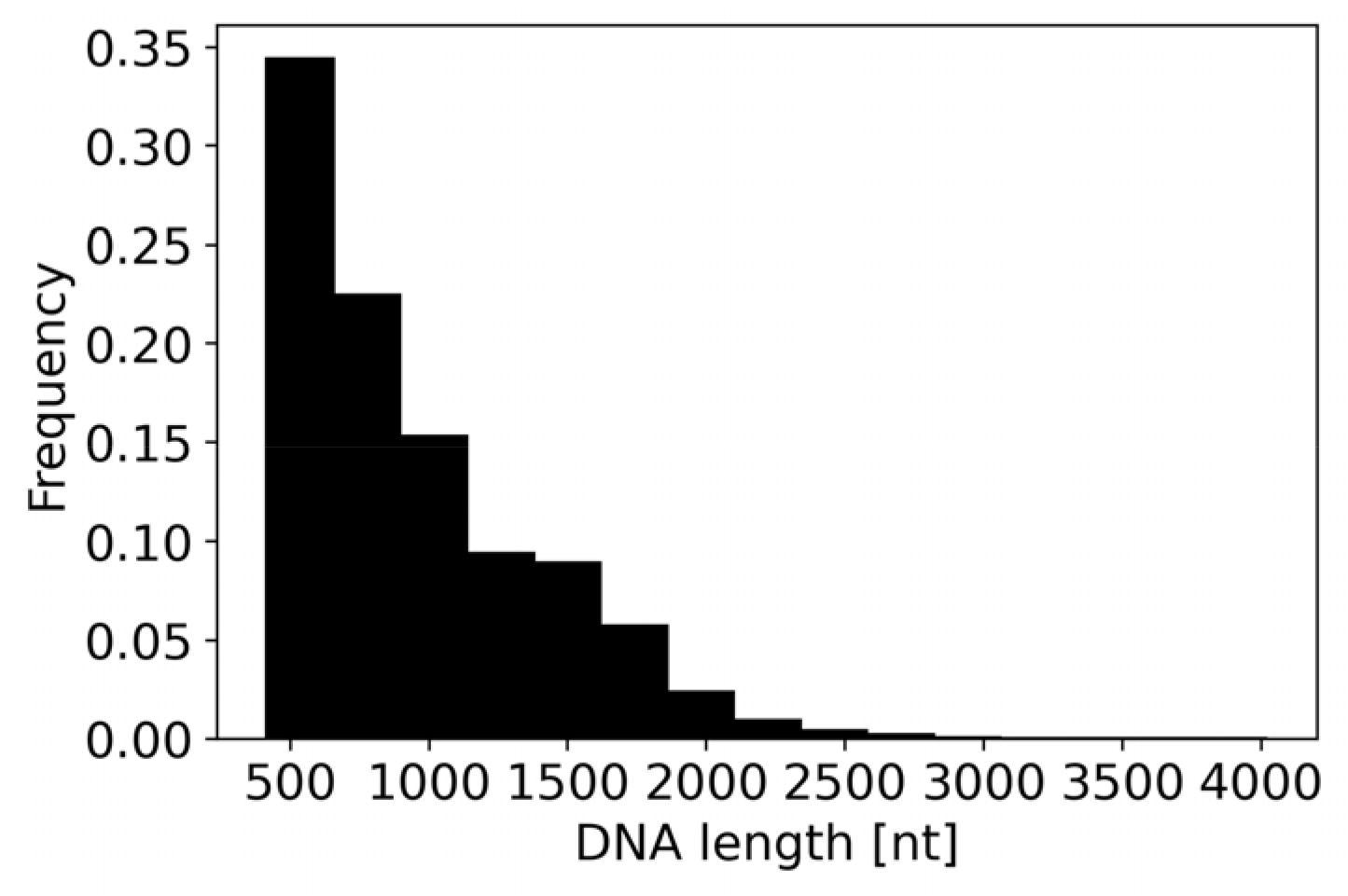
Distribution of lengths of DNA inserts for tagging or deleting S288C genes designed by PIPOline with default parameters. For tagging, the length of the insert does not include the tag nor the linker, which the user needs to choose (and ensure compatibility with restriction sites in the plasmid).

**Supplementary Figure 2.**
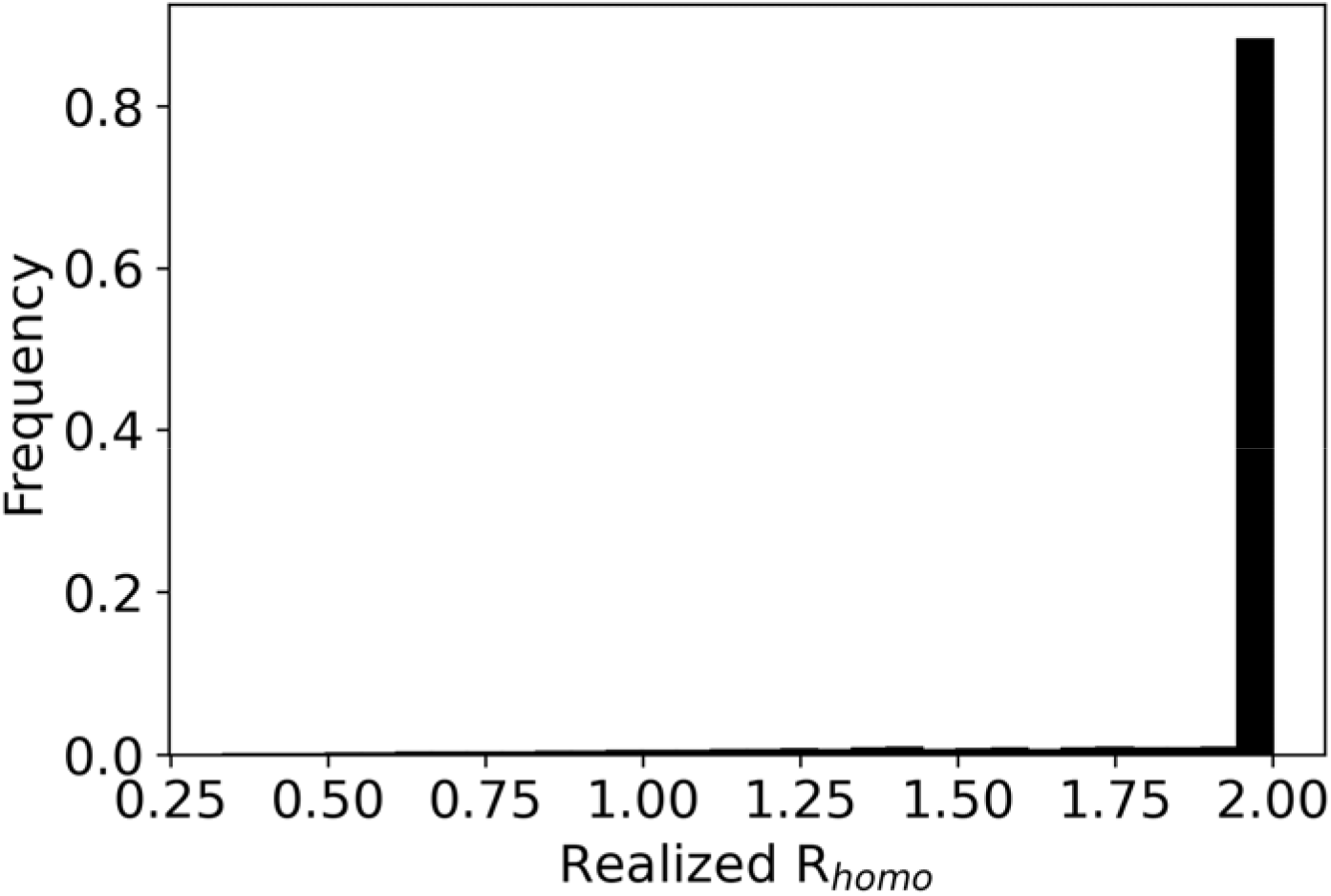
Distribution of realized R_homo_ parameters for tagging or deleting all S288C genes when using the default input value of 2 for R_homo_. The realized R_homo_ can be smaller than the desired R_homo_ if the rules for PIPOline cannot be satisfied otherwise. The average R_homo_ is (1.93 ± 0.22 STD).

Supplementary Note 1: List of PIPOline input parameters

| Argument | Description | Default Value |
| --- | --- | --- |
| --backbone_path | Plasmid backbone used for PIPO | './Test_examples/pETUL_backbone.fsa' |
| --MCS_start_ind | Starting nucleotide of the multiple cloning site sequence (1-based counting) | 1 |
| --MCS_end_ind | Ending nucleotide of the multiple cloning site sequence (1-based counting) | 108 |
| --min_homology | Minimal length of DNA sequence used for homologous recombination during pop-in | 70 |
| --R | Ratio between DNA sequence length used for pop-out vs. pop-in | 2 |
| --Gene_path | ORF of the gene with 1000 bp upstream and downstream (in FASTA format) | './Test_examples/CLB2_3p_labeling/CLB2_pm_1000.fsa' |
| --linker_path | Linker | './Test_examples/long_linker.fsa' |
| --modality | Modality of the program that can assume one of the following values:<br>- 0 for gene deletion<br>- 5 for gene tagging at the 5' terminus<br>- 3 for gene tagging at the 3' terminus | 0 |
| --enzyme_path | List of enzymes that can be used for cloning and plasmid linearization prior to transformation | './Test_examples/raw_enzyme_list.txt' |
| --FPG_paths | List of fluorescent protein coding genes that should be considered during plasmid design | ['./Test_examples/CLB2_3p_labeling/mCherry_FPG.fsa',<br>'./Test_examples/CLB2_3 |
|  |  | p_labeling/ymNeonGreen_FPG.fsa'<br>] |
| --popular_enzyme_path | List of enzymes to be considered for adding around linker and FPG | - |
| --assembled_plasmid_name | Name of the output FASTA file | - |

Supplementary Note 2: Example of PIPOline calls for gene deletion and gene tagging

Call used for gene deletion:

~~~
python.exe ./main.py
--backbone_path ‘./Test_examples/pETUL_backbone.fsa’
--MCS_start_ind 1
--MCS_end_ind 108
--min_homology 70
--R 2
--Gene_path ‘./Test_examples/GAL4_deletion/GAL4_pm_1000_gene.fsa’
--modality 0 --enzyme_path ‘./Test_examples/raw_enzyme_list.txt’
~~~

Call used for gene tagging:

~~~
python.exe ./main.py
--backbone_path ./Test_examples/pETUL_backbone.fsa
--MCS_start_ind 1
--MCS_end_ind 108
--min_homology 70
--R 2
--Gene_path ./Test_examples/CLB2_3p_labeling/CLB2_pm_1000.fsa
--linker_path ./Test_examples/long_linker.fsa
--modality 5 --enzyme_path ./Test_examples/raw_enzyme_list.txt
--popular_enzyme_path ./Test_examples/Popular_enzymes.txt
--FPG_paths ./Test_examples/CLB2_3p_labeling/mCherry_FPG.fsa
./Test_examples/CLB2_3p_labeling/ymNeonGreen_FPG.fsa
./Test_examples/CLB2_3p_labeling/ymTq2_FPG.fsa
~~~

Supplementary Note 3: Sequence of pETURALEU backbone

The following sequence is also supplied with annotations in gbk format in the Supplementary Files:

GGTACCGGGCCCCCCCTCGAGGTCGACGGTATCGATAAGCTTGATATCGAATTCCTGCAGCCCGGGGGATCCACTAGT TCTAGAGCGGCCGCCACCGCGGTGGAGCTCCAGCTTTTGTTCCCTTTAGTGAGGGTTAATTGCGCGCTTGGCGTAATC ATGGTCATAGCTGTTTCCTGTGTGAAATTGTTATCCGCTCACAATTCCACACAACATAcGAGCCGGAAGCATAAAGTGT AAAGCCTGGGGTGCCTAATGAGTGAGcTAACTCACATTAATTGCGTTGCGCTCACTGCCCGCTTTCCAGTCGGGAAAC CTGTCGTGCCAGCTGCATTAATGAATCGGCCAACGCGCGGGGAGAGGCGGTTTGCGTATTGGGCGCTCTTCCGCTTCC TCGCTCACTGACTCGCTGCGCTCGGTCGTTCGGCTGCGGCGAGCGGTATCAGCTCACTCAAAGGCGGTAATACGGTTA TCCACAGAATCAGGGGATAACGCAGGAAAGAACATGTGAGCAAAAGGCCAGCAAAAGGCCAGGAACCGTAAAAAG GCCGCGTTGCTGGCGTTTTTCCATAGGCTCCGCCCCCCTGACGAGCATCACAAAAATCGACGCTCAAGTCAGAGGTGG CGAAACCCGACAGGACTATAAAGATACCAGGCGTTTCCCCCTGGAAGCTCCCTCGTGCGCTCTCCTGTTCCGACCCTGC CGCTTACCGGATACCTGTCCGCCTTTCTCCCTTCGGGAAGCGTGGCGCTTTCTCATAGCTCACGCTGTAGGTATCTCAG TTCGGTGTAGGTCGTTCGCTCCAAGCTGGGCTGTGTGCACGAACCCCCCGTTCAGCCCGACCGCTGCGCCTTATCCGG TAACTATCGTCTTGAGTCCAACCCGGTAAGACACGACTTATCGCCACTGGCAGCAGCCACTGGTAACAGGATTAGCAG AGCGAGGTATGTAGGCGGTGCTACAGAGTTCTTGAAGTGGTGGCCTAACTACGGCTACACTAGAAGGACAGTATTTG GTATCTGCGCTCTGCTGAAGCCAGTTACCTTCGGAAAAAGAGTTGGTAGCTCTTGATCCGGCAAACAAACCACCGCTG GTAGCGGTGGTTTTTTTGTTTGCAAGCAGCAGATTACGCGCAGAAAAAAAGGATCTCAAGAAGATCCTTTGATCTTTT CTACGGGGTCTGACGCTCAGTGGAACGAAAACTCACGTTAAGGGATTTTGGTCATGAGATTATCAAAAAGGATCTTCA CCTAGATCCTTTTAAATTAAAAATGAAGTTTTAAATCAATCTAAAGTATATATGAGTAAACTTGGTCTGACAGTTACCA ATGCTTAATCAGTGAGGCACCTATCTCAGCGATCTGTCTATTTCGTTCATCCATAGTTGCCTGACTCCCCGTCGTGTAGA TAACTACGATACGGGAGGGCTTACCATCTGGCCCCAGTGCTGCAATGATACCGCGAGACCCACGCTCACCGGCTCCAG ATTTATCAGCAATAAACCAGCCAGCCGGAAGGGCCGAGCGCAGAAGTGGTCCTGCAACTTTATCCGCCTCCATCCAGT CTATTAATTGTTGCCGGGAAGCTAGAGTAAGTAGTTCGCCAGTTAATAGTTTGCGCAACGTTGTTGCCATTGCTACAG GCATCGTGGTGTCACGCTCGTCGTTTGGTATGGCTTCATTCAGCTCCGGTTCCCAACGATCAAGGCGAGTTACATGATC CCCCATGTTGTGCAAAAAAGCGGTTAGCTCCTTCGGTCCTCCGATCGTTGTCAGAAGTAAGTTGGCCGCAGTGTTATC ACTCATGGTTATGGCAGCACTGCATAATTCTCTTACTGTCATGCCATCCGTAAGATGCTTTTCTGTGACTGGTGAGTACT CAACCAAGTCATTCTGAGAATAGTGTATGCGGCGACCGAGTTGCTCTTGCCCGGCGTCAATACGGGATAATACCGCGC CACATAGCAGAACTTTAAAAGTGCTCATCATTGGAAAACGTTCTTCGGGGCGAAAACTCTCAAGGATCTTACCGCTGT TGAGATCCAGTTCGATGTAACCCACTCGTGCACCCAACTGATCTTCAGCATCTTTTACTTTCACCAGCGTTTCTGGGTGA GCAAAAACAGGAAGGCAAAATGCCGCAAAAAAGGGAATAAGGGCGACACGGAAATGTTGAATACTCATACTCTTCCT TTTTCAATATTATTGAAGCATTTATCAGGGTTATTGTCTCATGAGCGGATACATATTTGAATGTATTTAGAAAAATAAAC AAATAGGGGTTCCGCGCACATTTCCCCGAAAAGTGCCACCTGACGTCTAAGAAACCATTATTATCATGACATTAACCTA TAAAAATAGGCGTATCACGAGGCCCTTTCGTCTCGCGCGTTTCGGTGATGACGGTGAAAACCTCTGACACATGCAGCT CCcGGACTCTAATTTGTGAGTTTAGTATACATGCATTTACTTATAATACAGTTTTTTAGTTTTGCTGGCCGCATCTTCTCA AATATGCTTCCCAGCCTGCTTTTCTGTAACGTTCACCCTCTACCTTAGCATCCCTTCCCTTTGCAAATAGTCCTCTTCCAA CAATAATAATGTCAGATCCTGTAGAGACCACATCATCCACGGTTCTATACTGTTGACCCAATGCGTCTCCCTTGTCATCT AAACCCACACCGGGTGTCATAATCAACCAATCGTAACCTTCATCTCTTCCACCCATGTCTCTTTGAGCAATAAAGCCGAT AACAAAATCTTTGTCGCTCTTCGCAATGTCAACAGTACCCTTAGTATATTCTCCAGTAGATAGGGAGCCCTTGCATGAC AATTCTGCTAACATCAAAAGGCCTCTAGGTTCCTTTGTTACTTCTTCTGCCGCCTGCTTCAAACCGCTAACAATACCTGG GCCCACCACACCGTGTGCATTCGTAATGTCTGCCCATTCTGCTATTCTGTATACACCCGCAGAGTACTGCAATTTGACT GTATTACCAATGTCAGCAAATTTTCTGTCTTCGAAGAGTAAAAAATTGTACTTGGCGGATAATGCCTTTAGCGGCTTAA CTGTGCCCTCCATGGAAAAATCAGTCAAGATATCCACATGTGTTTTTAGTAAACAAATTTTGGGACCTAATGCTTCAAC TAACTCCAGTAATTCCTTGGTGGTACGAACATCCAATGAAGCACACAAGTTTGTTTGCTTTTCGTGCATGATATTAAAT AGCTTGGCAGCAACAGGACTAGGATGAGTAGCAGCACGTTCCTTATATGTAGCTTTCGACATGATTTATCTTCGTTTCC TGCAGGTTTTTGTTCTGTGCAGTTGGGTTAAGAATACTGGGCAATTTCATGTTTCTTCAACACTACATATGCGTATATAT ACCAATCTAAGTCTGTGCTCCTTCCTTCGTTCTTCCTTCTGTTCGGAGATTACCGAATCAAAAAAATTTCAAGGAAACCG AAATCAAAAAAAAGAATAAAAAAAAAATGATGAATTGAAAAGGTGGTATGGTGCACTCTCAGTACTCCgGGAGACGG TCACAGCTTGTCTGTAAGCGGATGCCGGGAGCAGACAAGCCCGTCAGGGCGCGTCAGCGGGTGTTGGCGGGTGTCG GGGCTGGCTTAACTATGCGGCATCAGAGCAGATTGTACTGAGAGTGCACCATATCGACTACGTCGTTAAGGCCGTTTC TGACAGAGTAAAATTCTTGAGGGAACTTTCACCATTATGGGAAATGGTTCAAGAAGGTATTGACTTAAACTCCATCAA ATGGTCAGGTCATTGAGTGTTTTTTATTTGTTGTATTTTTTTTTTTTTAGAGAAAATCCTCCAATATATAAATTAGGAATC ATAGTTTCATGATTTTCTGTTACACCTAACTTTTTGTGTGGTGCCCTCCTCCTTGTCAATATTAATGTTAAAGTGCAATTC TTTTTCCTTATCACGTTGAGCCATTAGTATCAATTTGCTTACCTGTATTCCTTTACATCCTCCTTTTTCTCCTTCTTGATAA ATGTATGTAGATTGCGTATATAGTTTCGTCTACCCTATGAACATATTCCATTTTGTAATTTCGTGTCGTTTCTATTATGAA TTTCATTTATAAAGTTTATGTACAAATATCATAAAAAAAGAGAATCTTTTTAAGCAAGGATTTTCTTAACTTCTTCGGCG ACAGCATCACCGACTTCGGTGGTACTGTTGGAACCACCTAAATCACCAGTTCTGATACCTGCATCCAAAACCTTTTTAA CTGCATCTTCAATGGCCTTACCTTCTTCAGGCAAGTTCAATGACAATTTCAACATCATTGCAGCAGACAAGATAGTGGC GATAGGGTCAACCTTATTCTTTGGCAAATCTGGAGCAGAACCGTGGCATGGTTCGTACAAACCAAATGCGGTGTTCTT GTCTGGCAAAGAGGCCAAGGACGCAGATGGCAACAAACCCAAGGAACCTGGGATAACGGAGGCTTCATCGGAGATG ATATCACCAAACATGTTGCTGGTGATTATAATACCATTTAGGTGGGTTGGGTTCTTAACTAGGATCATGGCGGCAGAA TCAATCAATTGATGTTGAACCTTCAATGTAGGGAATTCGTTCTTGATGGTTTCCTCCACAGTTTTTCTCCATAATCTTGA AGAGGCCAAAAGATTAGCTTTATCCAAGGACCAAATAGGCAATGGTGGCTCATGTTGTAGGGCCATGAAAGCGGCCA TTCTTGTGATTCTTTGCACTTCTGGAACGGTGTATTGTTCACTATCCCAAGCGACACCATCACCATCGTCTTCCTTTCTCT TACCAAAGTAAATACCTCCCACTAATTCTCTGACAACAACGAAGTCAGTACCTTTAGCAAATTGTGGCTTGATTGGAGA TAAGTCTAAAAGAGAGTCGGATGCAAAGTTACATGGTCTTAAGTTGGCGTACAATTGAAGTTCTTTACGGATTTTTAG TAAACCTTGTTCAGGTCTAACACTACCGGTACCCCATTTAGGACCACCCACAGCACCTAACAAAACGGCATCAGCCTTC TTGGAGGCTTCCAGCGCCTCATCTGGAAGTGGAACACCTGTAGCATCGATAGCAGCACCACCAATTAAATGATTTTCG AAATCGAACTTGACATTGGAACGAACATCAGAAATAGCTTTAAGAACCTTAATGGCTTCGGCTGTGATTTCTTGACCAA CGTGGTCACCTGGCAAAACGACGATCTTCTTAGGGGCAGACATTAGAATGGTATATCCTTGAAATATATATATATATAT TGCTGAAATGTAAAAGGTAAGAAAAGTTAGAAAGTAAGACGATTGCTAACCACCTATTGGAAAAAACAATAGGTCCT TAAATAATATTGTCAACTTCAAGTATTGTGATGCAAGCATTTAGTCATGAACGCTTCTCTATTCTATATGAAAAGCCGG TTCCGGCGCTCTCACCTTTCCTTTTTCTCCCAATTTTTCAGTTGAAAAAGGTATATGCGTCAGGCGACCTCTGAAATTAA CAAAAAATTTCCAGTCATCGAATTTGATTCTGTGCGATAGCGCCCCTGTGTGTTCTCGTTATGTTGAGGAAAAAAATAA TGGTTGCTAAGAGATTCGAACTCTTGCATCTTACGATACCTGAGTATTCCCACAGTTAACTGCGGTCAAGATATTTCTT GAATCAGGCGCCTTAGACCGCTCGGCCAAACAACCAATTACTTGTTGAGAAATAGAGTATAATTATCCTATAAATATA ACGTTTTTGAACACACATGAACAAGGAAGTACAGGACAATTGATTTTGAAGAGAATGTGGATTTTGATGTAATTGTTG GGATTCCATTTTTAATAAGGCAATAATATTAGGTATGTAGATATACTAGAAGTTCTCCTCGACCGGTCGATATGCGGTG TGAAATACCGCACAGATGCGTAAGGAGAAAATACCGCATCAGGAAATTGTAAgCGTTAATATTTTGTTAAAATTCGCG TTAAATTTTTGTTAAATCAGCTCATTTTTTAACCAATAGGCCGAAATCGGCAAAATCCCTTATAAATCAAAAGAATAGA CCGAGATAGGGTTGAGTGTTGTTCCAGTTTGGAACAAGAGTCCACTATTAAAGAACGTGGACTCCAACGTCAAAGGG CGAAAAACCGTCTATCAGGGCGATGGCCCACTACGTGAACCATCACCCTAATCAAGTTTTTTGGGGTCGAGGTGCCGT AAAGCACTAAATCGGAACCCTAAAGGGAGCCCCCGATTTAGAGCTTGACGGGGAAAGCCGGCGAACGTGGCGAGAA AGGAAGGGAAGAAAGCGAAAGGAGCGGGCGCTAGGGCGCTGGCAAGTGTAGCGGTCACGCTGCGCGTAACCACCA CACCCGCCGCGCTTAATGCGCCGCTACAGGGCGCGTCCATTCGCCATTCAGGCTGCGCAACTGTTGGGAAGGGCGAT CGGTGCGGGCCTCTTCGCTATTACGCCAGCTGGCGAAAGGGGGATGTGCTGCAAGGCGATTAAGTTGGGTAACGCCA GGGTTTTCCCAGTCACGACGTTGTAAAACGACGGCCAGTGAGCGCGCGTAATACGACTCACTATAGGGCGAATTG

Supplementary Note 4: Restriction enzymes and their cut sites

The following default list of restriction enzymes and cut sites is included in the PIPOline package in file raw_enzyme_list.txt:

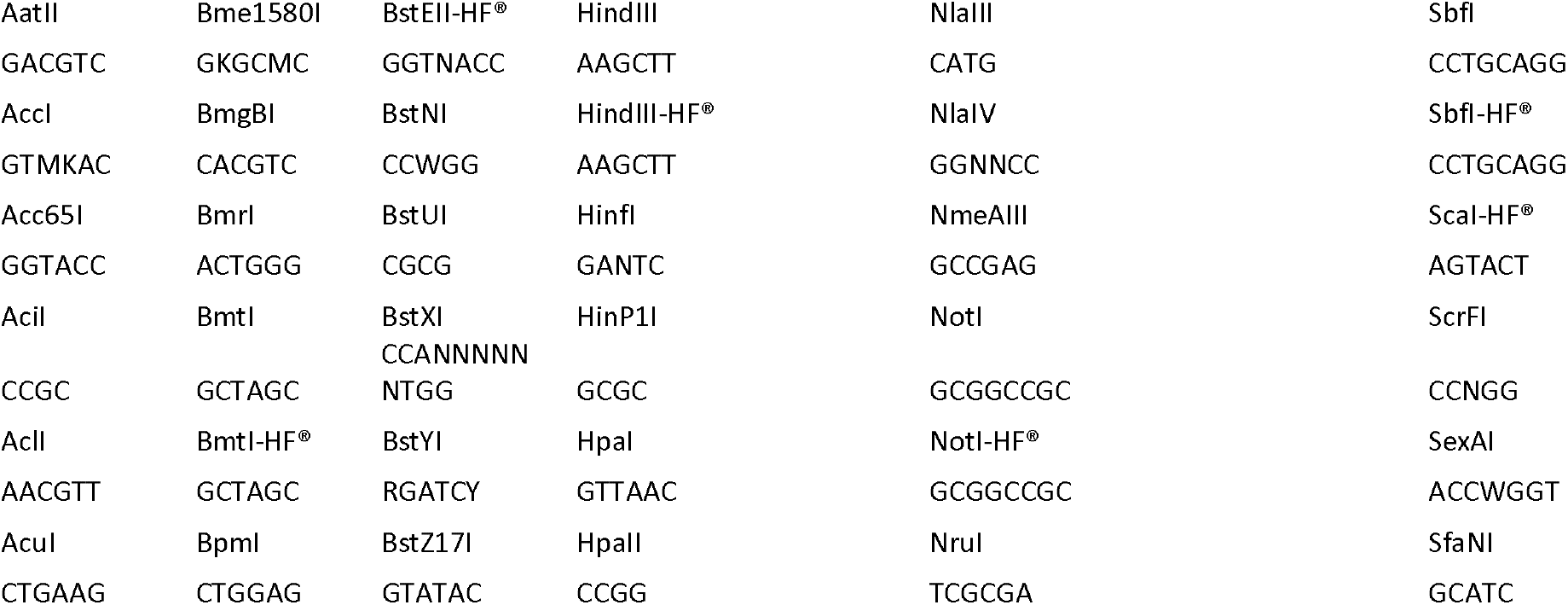
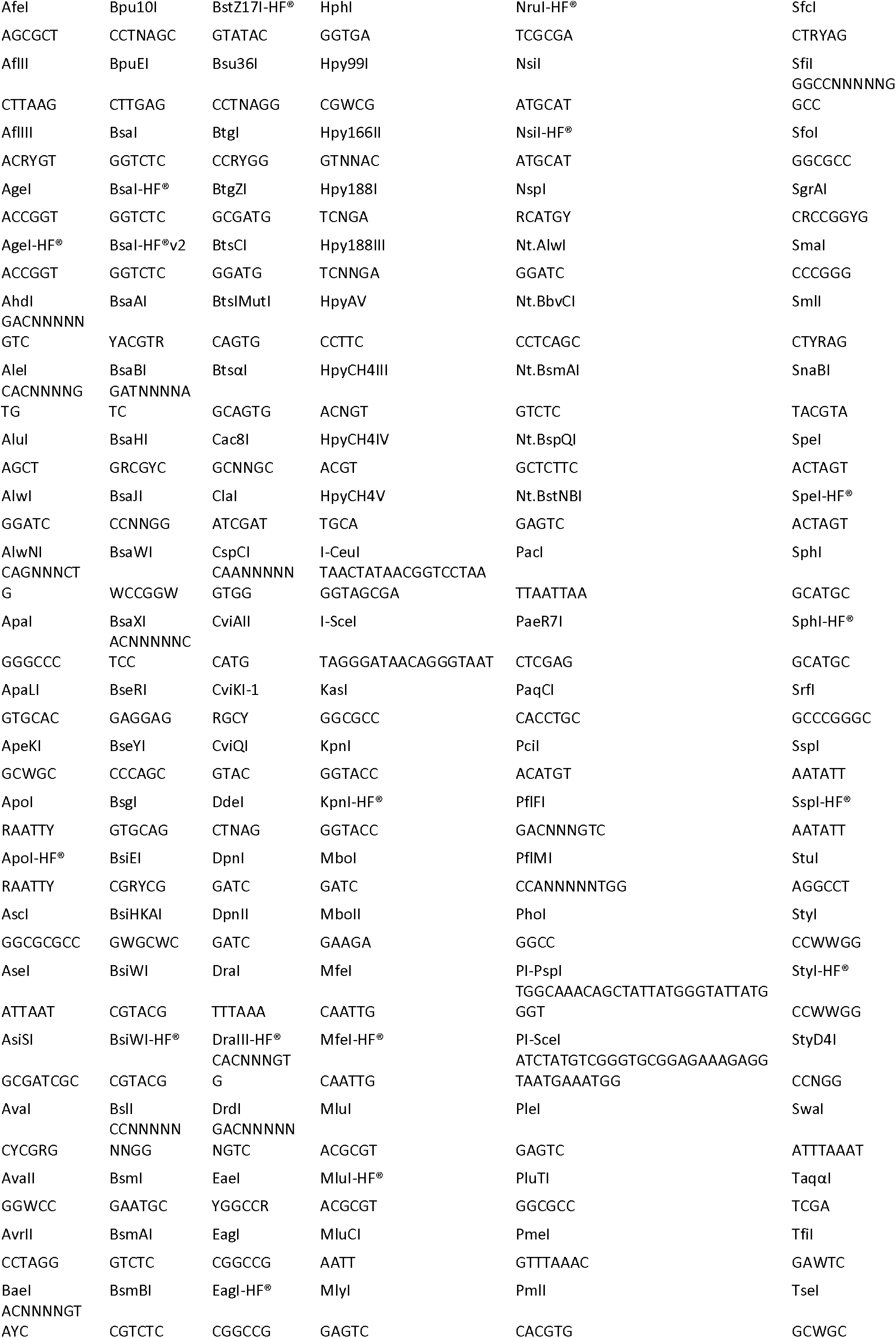
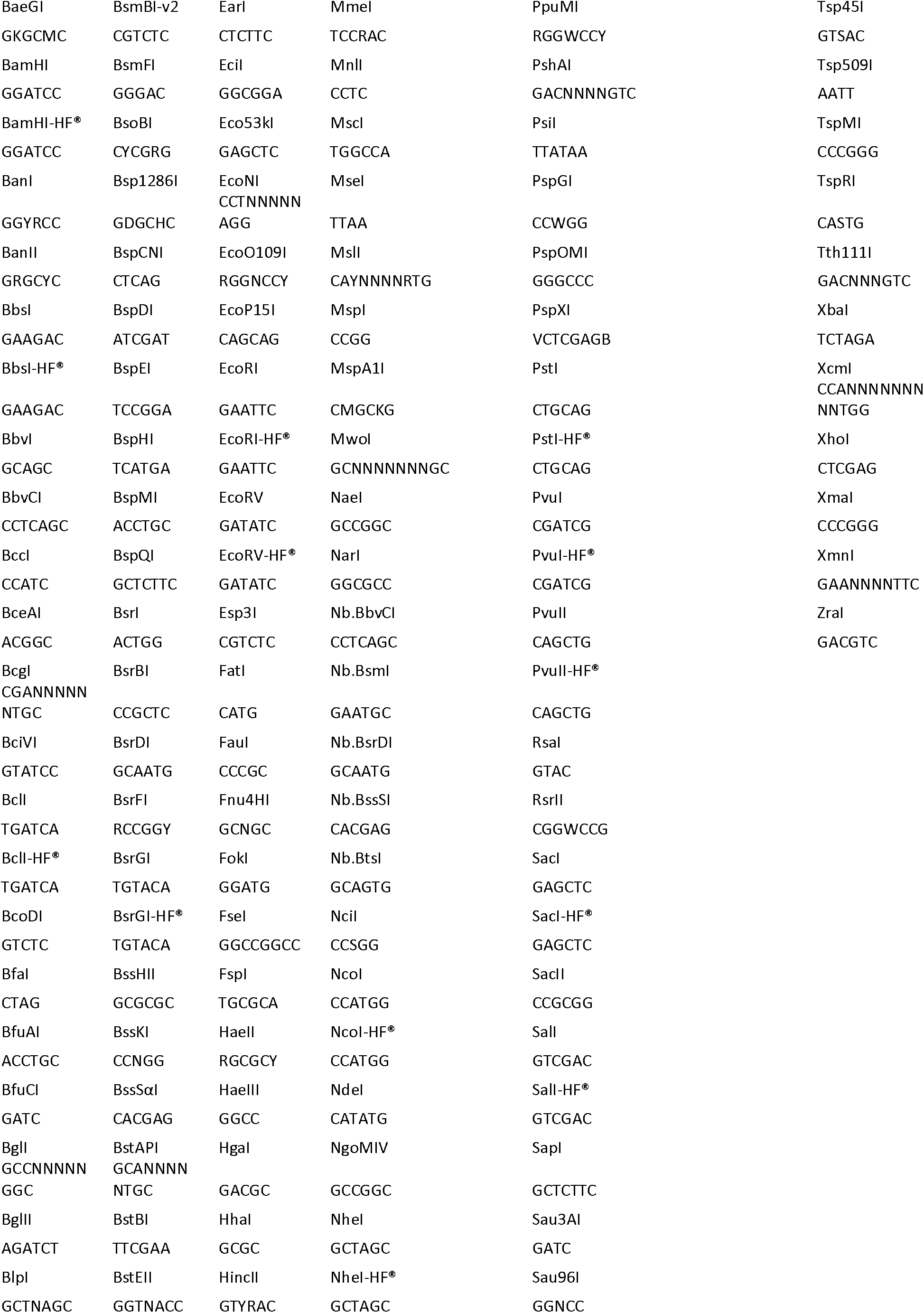

Supplementary Note 5: Default linker used for tagging

CCGCGGGGTGCTTCTGTTGGTGCTTCTGTTTCTGTTGGTCCGCGGTGGAGCTCAATG

Supplementary Note 6: Sequence of yeast-optimized mScarletI (ymScarletI) from ref. ^35^ with restriction sites for frequently-used restriction enzymes removed

The following sequence is also supplied with annotations in gbk format in the Supplementary Files:

ATGGTTAGTAAAGGTGAAGCTGTTATAAAAGAATTTATGAGGTTTAAAGTTCATATGGAAGGTTCAATGAATGGTCAT GAATTTGAAATTGAAGGTGAAGGTGAAGGTAGACCATATGAAGGTACACAAACTGCTAAATTGAAAGTTACTAAAGG TGGTCCATTACCATTTTCTTGGGATATTTTGTCTCCACAATTCATGTATGGTTCAAGAGCTTTTATTAAGCATCCTGCTG ATATTCCAGATTATTATAAACAATCTTTTCCTGAAGGTTTTAAATGGGAAAGAGTTATGAATTTTGAAGATGGTGGTGC TGTTACTGTTACACAAGATACTTCTTTGGAAGATGGTACTTTAATCTATAAAGTTAAATTGAGAGGTACTAATTTTCCAC CAGACGGTCCAGTTATGCAAAAGAAAACTATGGGTTGGGAAGCATCTACTGAAAGATTGTATCCTGAAGATGGTGTT TTGAAAGGAGACATTAAGATGGCTTTGAGATTGAAAGATGGTGGTAGATACCTGGCCGACTTCAAGACCACCTATAA AGCTAAAAAACCAGTTCAAATGCCAGGTGCATATAATGTTGATAGAAAGTTAGATATAACGTCGCATAACGAGGACTA TACTGTAGTTGAACAATATGAACGTAGTGAAGGTAGACATAGTACCGGAGGAATGGATGAATTGTATAAATAA

**Supplementary Table 1:** Strains used in the study.

| Strain name | Genotype | Source |
| --- | --- | --- |
| yVG2873 | *W303 MATa WHI5::pVG264::URA3, LEU2* | This study |
| yVG2874 | *W303 MATa WHI5::pVG265::URA3, LEU2* | This study |
| yVG2875 | *W303 MATa WHI5::pVG266::URA3, LEU2* | This study |
| yVG2876 | *W303 MATa WHI5::pVG267::URA3, LEU2* | This study |
| yVG2877 | *W303 MATa WHI5::pVG268::URA3, LEU2* | This study |
| yVG2878 | *W303 MATa HTB2::pVG288::URA3, LEU2* | This study |
| yVG2879 | *W303 MATa HTB2::pVG289::URA3, LEU2* | This study |
| yVG2880 | *W303 MATa HTB2::pVG290::URA3, LEU2* | This study |
| yVG2881 | *W303 MATa HTB2::pVG291::URA3, LEU2* | This study |

**Supplementary Table 2:** Plasmids used in the study. The annotated sequences for the plasmids below are included as gbk files in the Supplementary Files:

| Plasmid name | Description | Yeast selection marker | Bacterial selection marker | Source of the plasmid |
| --- | --- | --- | --- | --- |
| pETURALEU | Empty backbone containing both *URA3* and *LEU2* selectable markers | *URA3, LEU2* | *Amp* | This study |
| pVG254 | Whi5 marking plasmid using ymScarletI and $R_{\text{homo}} = 0.5$ | *URA3, LEU2* | *Amp* | This study |
| pVG265 | Whi5 marking plasmid using ymScarletI and $R_{\text{homo}} = 1$ | *URA3, LEU2* | *Amp* | This study |
| pVG266 | Whi5 marking plasmid using ymScarletI and $R_{\text{homo}} = 1.5$ | *URA3, LEU2* | *Amp* | This study |
| pVG267 | Whi5 marking plasmid using ymScarletI and $R_{\text{homo}} = 2$ | *URA3, LEU2* | *Amp* | This study |
| pVG268 | Whi5 marking plasmid using ymScarletI with $R_{\text{homo}} = 4$ | *URA3, LEU2* | *Amp* | This study |
| pVG288 | Htb2 marking plasmid using ymScarletI and $R_{\text{homo}} = 0.5$ | *URA3, LEU2* | *Amp* | This study |
| pVG289 | Htb2 marking plasmid using ymScarletI and $R_{\text{homo}} = 1$ | *URA3, LEU2* | *Amp* | This study |
| pVG290 | Htb2 marking plasmid using ymScarletI and $R_{\text{homo}} = 1.5$ | *URA3, LEU2* | *Amp* | This study |
| pVG291 | Htb2 marking plasmid using ymScarletI and $R_{\text{homo}} = 2$ | *URA3, LEU2* | *Amp* | This study |

**Supplementary Table 3:**
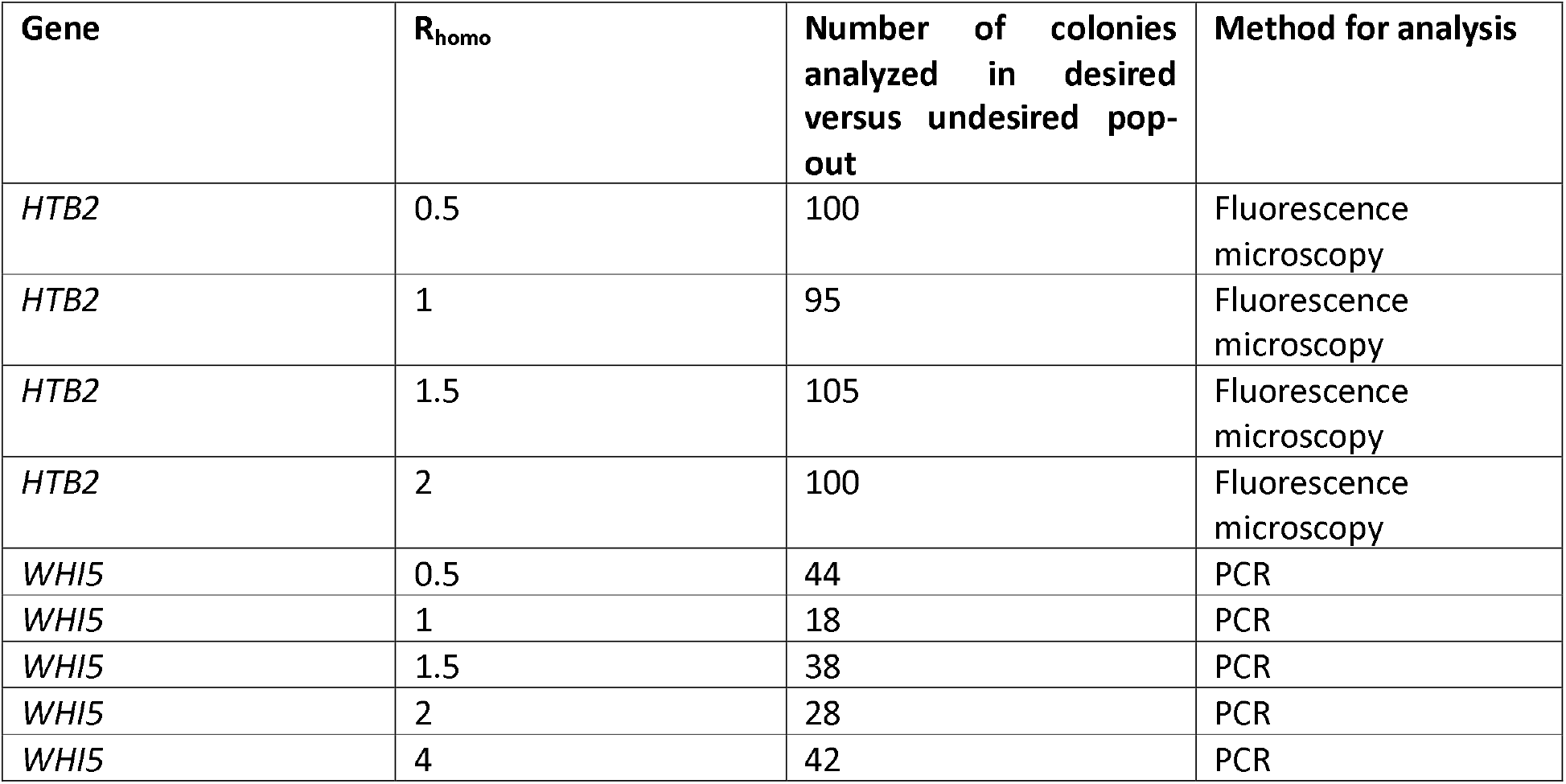
Number of analyzed colonies for Fig. 4.

| Gene | $R_{\text{homo}}$ | Number of colonies analyzed in desired versus undesired pop-out | Method for analysis |
| --- | --- | --- | --- |
| *HTB2* | 0.5 | 100 | Fluorescence microscopy |
| *HTB2* | 1 | 95 | Fluorescence microscopy |
| *HTB2* | 1.5 | 105 | Fluorescence microscopy |
| *HTB2* | 2 | 100 | Fluorescence microscopy |
| *WHI5* | 0.5 | 44 | PCR |
| *WHI5* | 1 | 18 | PCR |
| *WHI5* | 1.5 | 38 | PCR |
| *WHI5* | 2 | 28 | PCR |
| *WHI5* | 4 | 42 | PCR |

